# Size-structured mating niche differentiation in the male-polyphenic mite *Rhizoglyphus robini*

**DOI:** 10.1101/2021.12.24.474119

**Authors:** Flor T. Rhebergen, Maarten Wijstma, Isabel M. Smallegange

## Abstract

Condition-dependent expression of alternative male morphs evolves when males of different sizes experience different mating niches, requiring different morphologies. Such mating niche differentiation can be due to competitive asymmetry between large and small males in contests over mating opportunities. Here, we tested the hypothesis that aggressive interactions among males cause size-structured mating niches in an acarid mite with condition-dependent male polyphenism: the bulb mite *Rhizoglyphus robini*. In this species, large males mature as armed fighters with enlarged legs, and small males as scramblers without modified legs. We staged experimental dyadic contests over a mating opportunity between either a pair of fighter males, or a fighter and a scrambler male. We predicted that the larger male would have a higher likelihood of mating first in contests among fighters, that the fighter male would have a higher likelihood of mating first in fighter-scrambler contests, that fighters would have a higher likelihood of interrupting ongoing copulations if they are larger than their mating rival, and that copulations in the presence of a fighter rival therefore last shorter than copulations with a scrambler rival present. We found that in contests among fighters, the larger male had a higher probability of mating first. In contests among a fighter and scrambler, the fighter male was more likely to mate first regardless of the body size difference between the contestants. Ongoing copulations were only rarely interrupted by the rival male (always by a fighter), and the probability of interruption did not depend on the body size difference between the mating male and its rival. Copulations lasted shorter in the presence of a rival fighter, but this effect was not attributable to interruption of copulations. We conclude that the fighting niche is particularly accessible for large males, as larger males have a higher probability of winning pre-copulatory contests. Such mating niche differentiation likely contributes to evolutionary maintenance of condition-dependent male polyphenism, where small males are forced to adopt an alternative mating tactic and hence develop a dedicated morphology.

## Introduction

Male secondary sexual traits are often conspicuously variable within animal populations, with small males typically developing proportionally smaller or less exaggerated traits than larger males (Emlen and Nijhout 2000; Kodric-Brown et al. 2006; Lavine et al. 2015). In several arthropods, such condition-dependent expression of sexual traits is represented by discrete alternative male phenotypes (Danforth and Desjardins 1999; Tomkins 1999; Emlen et al. 2005; Cook and Bean 2006; Radwan 2009; Buzatto and Machado 2014; McCullough et al. 2015; Karplus and Barki 2019). It is a longstanding hypothesis that such condition-dependent polyphenism in males is maintained by social-status-dependent sexual selection for alternative mating tactics in small and large individuals (Dawkins 1980; Eberhard 1982; Parker 1982; Gross 1996; Tomkins and Hazel 2007). According to this idea, small males are physically weaker than large males, unlikely to compete successfully over mating opportunities in direct intrasexual competition with large males, and adaptively ‘make the best of a bad job’ by developing a morphology adapted for alternative, contest-avoiding mating behaviour such as sneaking (Gross 1996; Brockmann 2001; Taborsky et al. 2008). Hence, condition-dependent expression of alternative male morphologies is hypothesized to be maintained by the existence of different mating niches (*sensu* Shuster and Wade (2003)) for males of different sizes.

Evidence that condition-dependent polyphenism in male sexual traits evolves under disruptive selection due to size-structured mating niches is sparse, despite the relatively widespread occurrence of such polyphenisms (Dominey 1984; Gross 1996; Oliveira et al. 2008). Within arthropods, the evidence is largely confined to horned beetles in which small males develop a ‘minor’ phenotype without large horns. In these cases, fights among males over mating opportunities are generally won by the larger male, so that small males resort to fight-avoiding sneaking behaviour to gain access to a female (Emlen 1997; Moczek and Emlen 2000; Hongo 2007; McCullough and Simmons 2016; Mitchem et al. 2019). In contrast, tests for size-structured mating niches in male-diphenic earwigs (Styrsky and Van Rhein 1999; Forslund 2003), prawns (Barki et al. 1997), harvestmen (Buzatto and Machado 2008) and acarid mites (Michalczyk et al. 2018) have been inconclusive. Thus, while the evolution of male polyphenism can be explained by the existence of distinct and size-dependent mating niches in horned beetles, it remains poorly understood in other arthropods.

Here, we test the hypothesis that aggressive interactions among males cause size-structured mating niches in an acarid mite with condition-dependent polyphenism in a male secondary sexual trait: the bulb mite *Rhizoglyphus robini*. Confirmation of this hypothesis would strengthen the general idea that condition-dependent polyphenism in male sexual traits evolves under disruptive sexual selection due to size-structured mating niches. In bulb mites, large well-fed males develop a strongly enlarged third leg pair with sharp claws (the ‘fighter’ phenotype) (Radwan 1995; Smallegange 2011), which function as weapons in fights with rival males (Radwan and Klimas 2001; Radwan 2009). In contrast, small and poorly-fed males tend to develop an unmodified third leg pair, which is morphologically similar to the other legs (the ‘scrambler’ phenotype) (Radwan 1995; Smallegange 2011). Fighter males are more aggressive than scrambler males (Radwan et al. 2000), and can monopolize females by killing rivals (Radwan and Klimas 2001), while scramblers have a mobility advantage (Tomkins et al. 2011) and can obtain higher fertilization success from a single copulation (Van den Beuken et al. 2019). Male morph expression depends on body size immediately prior to metamorphosis, with larger males becoming fighters and smaller males scramblers (Radwan 1995; Smallegange 2011), although adult fighter males are on average not larger than adult scrambler males (Smallegange et al. 2012); fighters shrink during metamorphosis while scramblers do not (Radwan et al. 2002). In several closely related acarid mite species, fighter expression is additionally suppressed by pheromones signaling high colony density (Radwan 1995, 2001; Michalczyk et al. 2018); this does not happen in *R. robini* (Radwan 1995). Copulating male *R. robini* have been demonstrated to be vulnerable to attacks from rivals, and the presence of fighter males prevents the evolution of prolonged mate guarding, where pairs remain in copula after sperm transfer (Radwan and Siva-Jothy 1996; Skwierzyńska et al. 2018). Thus, the male fighting niche can be size-dependent in two different ways: the probability of eliminating a rival before copulation may depend on the size difference between the rivals, or the probability of successfully disrupting rival copulations may be size-dependent.

From our hypothesis that the fighting niche is size-dependent, we derive the following predictions. Firstly, in fights among fighters over a mating opportunity, the larger fighter male should have a higher probability of mating first than the smaller male (i.e. only large males occupy the fighting niche successfully; *prediction 1*). Secondly, in fights between a fighter and a scrambler over a mating opportunity, the fighter male should be more likely to mate first regardless of its size (i.e. scramblers are not well adapted to the fighting niche; *prediction 2*). Thirdly, ongoing copulations are disrupted by rival fighters, not by rival scramblers, and fighters have a higher probability of dislodging their mating rival if they are larger than their rival (*prediction 3*). Therefore, copulations should last longer if rivals are scramblers than if rivals are fighters (*prediction 4*). To test these predictions, we staged experimental dyadic male contests for a mating opportunity (a virgin female) with fighter-fighter and fighter-scrambler dyads, noted body size of both contestants, and scored which male mated first, whether the copulation was subsequently disrupted by the other male, and how long mating lasted.

## Materials and Methods

### Study species and experimental procedures

*Rhizoglyphus robini* occurs naturally on underground plant structures, where it can form dense populations that feed on fungi and rotting plant material (Díaz et al. 2000). Our *R. robini* stocks come from flower bulbs in storage rooms in Noord-Holland, The Netherlands, and were collected in December 2010. The stocks are kept in an unlit climate cabinet (25 °C; >90% humidity), in plastic containers (10 × 10 × 2.5 cm) with a plaster of Paris substrate, and are kept on a diet of baker’s yeast. The stock populations each contain tens of thousands of mites, and are mixed every six months to maintain genetic homogeneity. Waste is removed and water and food are added twice a week. In these circumstances, generation time (egg-to-egg) averages 11 days. *R. robini* has three juvenile stages (larva, protonymph, tritonymph), interspersed with quiescent phases during which individuals do not move or eat, but undergo incomplete metamorphosis to enter the next stage. Alternative male morphs (fighter or scrambler) are expressed only in the adult stage. In our stock populations, approximately 70-80% of the adult males are fighters.

We set up experimental contests between two males over a mating opportunity (a virgin female). The males in each contest were either both fighters, or a fighter and a scrambler. These males were randomly collected from the stocks, and then individually isolated in plastic tubes (height 5 cm, diameter 1.5 cm) with a moistened plaster substrate, containing *ad lib* food (yeast). The tubes were sealed with fine mesh, kept in place by a plastic lid and placed in an unlit climate cabinet (25 °C; >90% humidity). Because we wanted to distinguish the males individually in the experimental contests, we dyed the food of half of the males green, and the food of other half red, using food dye. As adult mites are slightly translucent, the colored food was visible inside of the digestive tract, which made them easily distinguishable. Since the males were collected as adults, and had likely mated before, we kept them isolated for 3-5 days before experimentation, which we assumed would build up motivation to mate. We obtained virgin females by randomly collecting quiescent tritonymphs from the stocks, and isolating them in plastic tubes (see above) containing *ad lib* food (yeast, undyed). The females emerged as virgin adults the next day, and were used in the experiments 3-5 days later.

The experimental contests took place in empty plastic tubes (height 5 cm, diameter 1.5 cm) with a moistened plaster substrate (hereafter: arena), positioned under a microscope. For each contest, we randomly picked a green-coloured and a red-coloured male from the males that had been isolated 3-5 days before. Before releasing the males in the arena, we photographed them ventrally to measure their body size (width of the idiosoma, at its broadest point) to the nearest μm, using a Zeiss Axiocam 105 color camera mounted on a Zeiss Stemi 200-C stereomicroscope, and ZEN 2 (Blue edition) software. After photographing them, we released the males in the arena, and left them to acclimate for ten minutes. We then added a virgin female to the arena, introduced at equal distance of both males. *Rhizoglyphus* males move rather slowly, and pairs have previously been shown to remain in copula from several minutes up to multiple hours, with a median of 20 minutes (Radwan and Siva-Jothy 1996; Skwierzyńska et al. 2018). Therefore, we monitored the arenas for approximately 10 seconds every minute. This allowed us to run multiple trials simultaneously, while being frequent enough to record which male mated first, the duration of copulation (to the nearest minute), and whether the non-copulating male was attempting to disrupt its rival’s copulation. Experimental contests would last until one male had finished copulating with the female, and were aborted if no copulation had occurred within an hour. Our sample size was largely determined by logistical constraints; we managed to stage 57 fighter-fighter contests, and 14 fighter-scrambler contests. In fighter-scrambler contests, males were easily distinguishable by phenotype, but were alternately dyed green or red anyway, to avoid confounding between contest type and the dying procedure.

### Statistics

Our first prediction was that in contests between fighters of different sizes, the larger fighter has a higher probability of winning contests and mating first. To test this prediction, we constructed a logistic regression model (generalized linear models with logit link function), with the difference in size between the green (fighter 1) and the red fighter (fighter 2) as explanatory variable (i.e. size of the green fighter minus size of the red fighter); the response variable was whether the green male (fighter 1) mated first or not. If the size difference between the fighters does not explain which fighter mates first, then the probability of the green (or red) fighter winning should be 0.5 (since males were randomly assigned to green or red coloured food), and there should not be a significant effect of the size difference between the fighters on the probability that the green fighter mated first.

Our second prediction was that in contests between a fighter and a scrambler, the fighter has a higher probability of winning irrespective of the body size difference. To test this prediction, we constructed a logistic regression model with the difference in size between the fighter and the scrambler as explanatory variable (i.e. the size of the fighter minus the size of the scrambler); the response variable was whether the fighter mated first or not. If the outcome of the fighter-scrambler contest is explained by the body size difference rather than male morph, then there should be a significant effect of the size difference between the fighter and the scrambler on the probability that the fighter mated first, and the probability that the fighter mates first should be 0.5 at zero body size difference. On the other hand, if the outcome of the fighter-scrambler contest is explained by male morph, then there should not be a significant effect of the size difference between the fighter and the scrambler, and the probability that the fighter mates first should overall be higher than 0.5.

Our third prediction was that in contests between fighters of different sizes, a fighter has a higher probability of dislodging its mating rival if it is larger than its rival. To test this prediction, we constructed a logistic regression model with the size difference between the mated and the unmated fighter as the explanatory variable (i.e. the size of the mated male minus the size of the unmated male); the response variable was whether the unmated fighter disrupted the copulation (yes or no). In all logistic regression models, we tested significance of explanatory variables using likelihood ratio tests (LRT).

Our fourth prediction was that copulations in the presence of a rival last longer if the rival is a scrambler rather than a fighter. We tested this prediction using a Kruskal-Wallis test, with as explanatory variable ‘mating type’ (i.e. either mated fighter with unmated fighter, or mated fighter with unmated scrambler; only three contests involved a mated scrambler with an unmated fighter, and were excluded from this analysis), and mating duration as response variable. For this analysis we could not use a parametric test, as the residuals were non-normally distributed even after log-transformation of the response variable.

All analyses were done in R v. 3.4.4 (R Core Team 2018), including packages *stats, dplyr* (Wickham et al. 2018) and *ggplot2* (Wickham et al. 2019).

## Results

Our first prediction was that in contests between fighters of different sizes, the larger fighter has a higher probability of winning contests and mating first. In dyadic contests for a mating opportunity among fighter males, the probability of mating first depended significantly on the size difference between the males, so that the larger fighter male had a higher probability of mating than the smaller male (LRT: χ^2^ = 7.54, df = 1, P = 0.006; Fig. 2). The estimated probability that the green-dyed male (male 1) mated first at zero size difference was 0.61 (95% CI: [0.47 – 0.73]), not significantly different from 0.5 (z = 1.52; P = 0.128).

**Figure 1.**
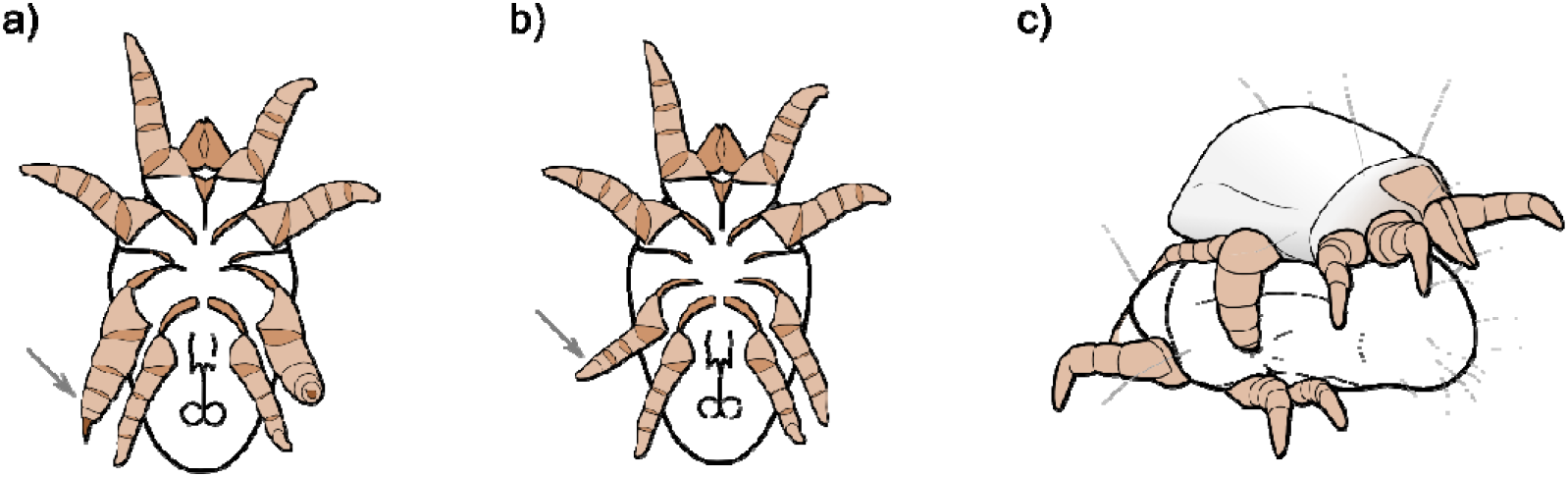
*Rhizoglyphus robini* alternative adult male phenotypes. Large juveniles become fighters (a, ventral view, head above), relatively small juveniles become scramblers (b, ventral view, head above). Note the difference in third leg morphology (arrows). Fighters use their enlarged legs with sharp tarsal claws as weapons in intrasexual fights (c).

**Figure 2.**
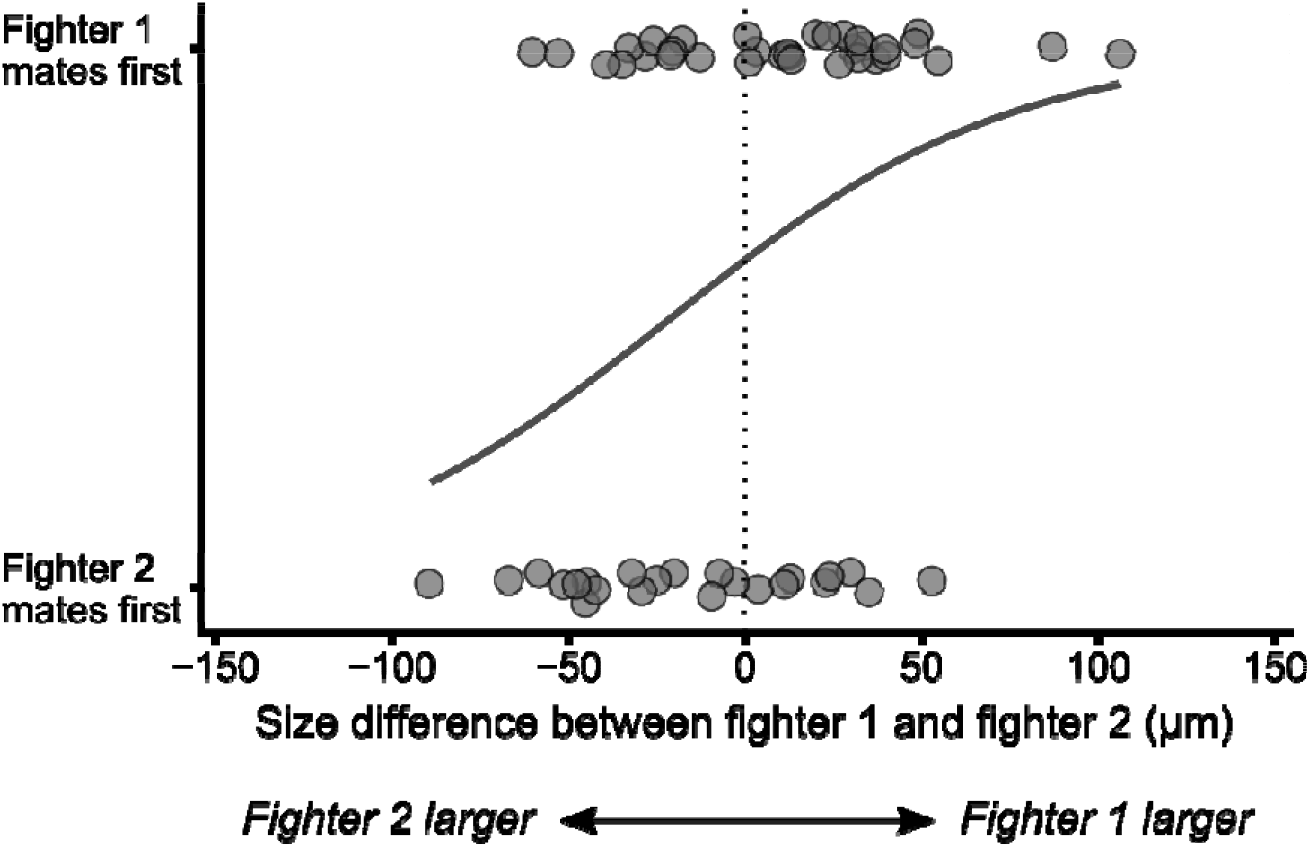
In fighter-fighter contests, the probability that a male mates first depends on its body size relative to its rival’s body size. Fighter 1 (the green male in experimental contests) has a higher probability of mating first if it is larger than fighter 2 (the red male in experimental contests), and vice versa, so the larger male has a higher probability of mating first. The black line represents the modelled probability of winning the contest and mating first. The dotted vertical line denotes equal body size, at which the probability that fighter 1 (the green male) mates first is estimated at 0.61 (95% CI: [0.47 – 0.73]).

Our second prediction was that in contests between a fighter and a scrambler, the fighter has a higher probability of winning irrespective of the body size difference. In dyadic contests between a fighter and a scrambler, there was no evidence that the size difference between the fighter and the scrambler contributed to which male mated first (LRT: χ^2^ = 0.00, df = 1, P = 0.982; Fig. 3). Instead, the probability of mating first was overall significantly higher for fighters (0.79, 95% CI: [0.50 – 0.93]; LRT: χ^2^ = 4.86, df = 1, P = 0.027; Fig. 3), regardless of the size difference.

**Figure 3.**
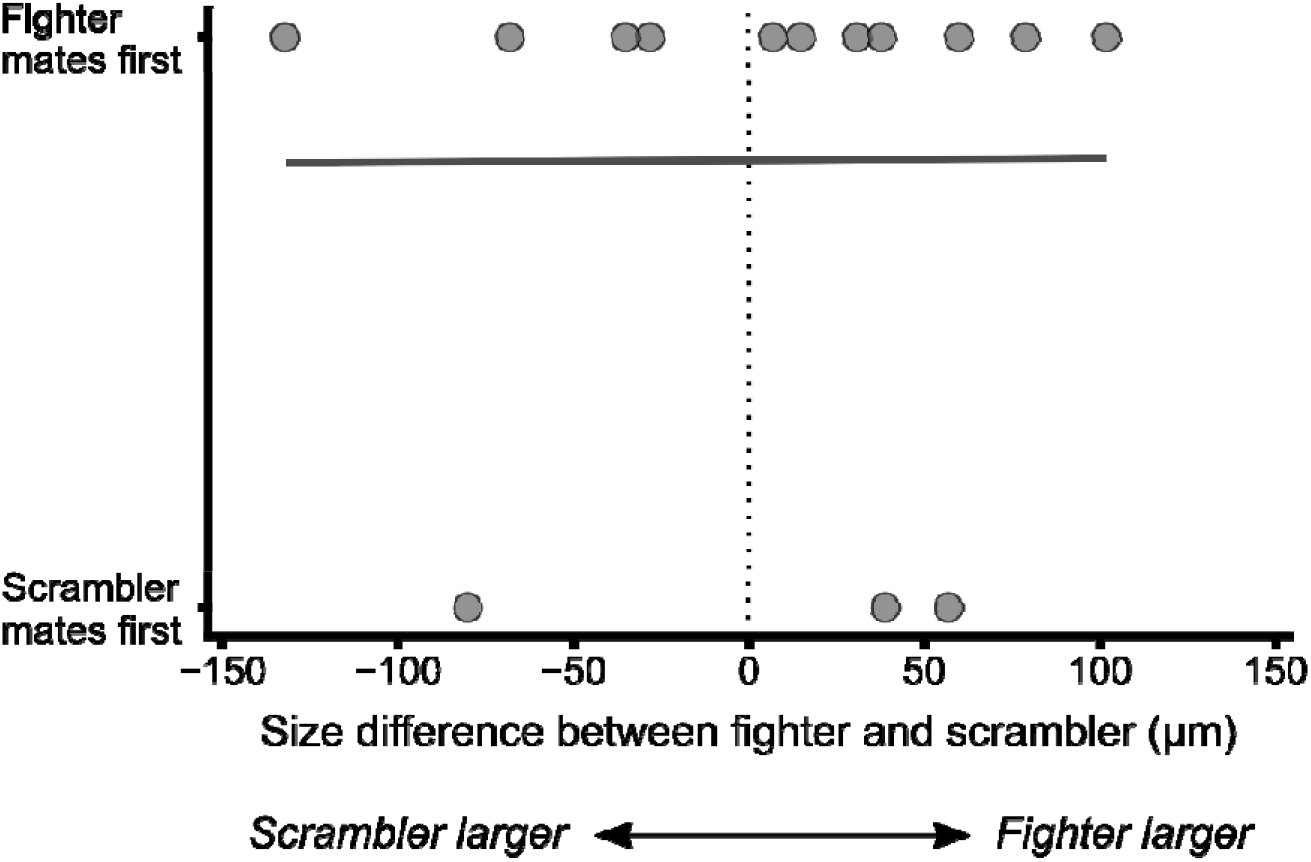
In fighter-scrambler contests, the probability that a male mates first does not depend on relative body size. Instead, fighters have an overall higher probability of winning the contest and mating first. The black line represents the modelled probability of winning the contest and mating first. The dotted vertical line denotes equal body size, at which the probability that the fighter mates first is estimated at 0.79 (95% CI: [0.50 – 0.93]).

Our third prediction was that in contests between fighters of different sizes, a fighter has a higher probability of dislodging its mating rival if it is larger than its rival. Copulations by fighters were only rarely interrupted by the rival that did not mate first (n=7; probability of interruption estimated at 0.12 (95% CI: [0.06 – 0.25]), and only in fighter-fighter contests. There was no evidence that the probability of such interruptions depended on the body size difference between the interrupting male and the mating male (LRT: χ^2^ = 0.00, df = 1, P = 0.992; Fig. 4).

**Figure 4.**
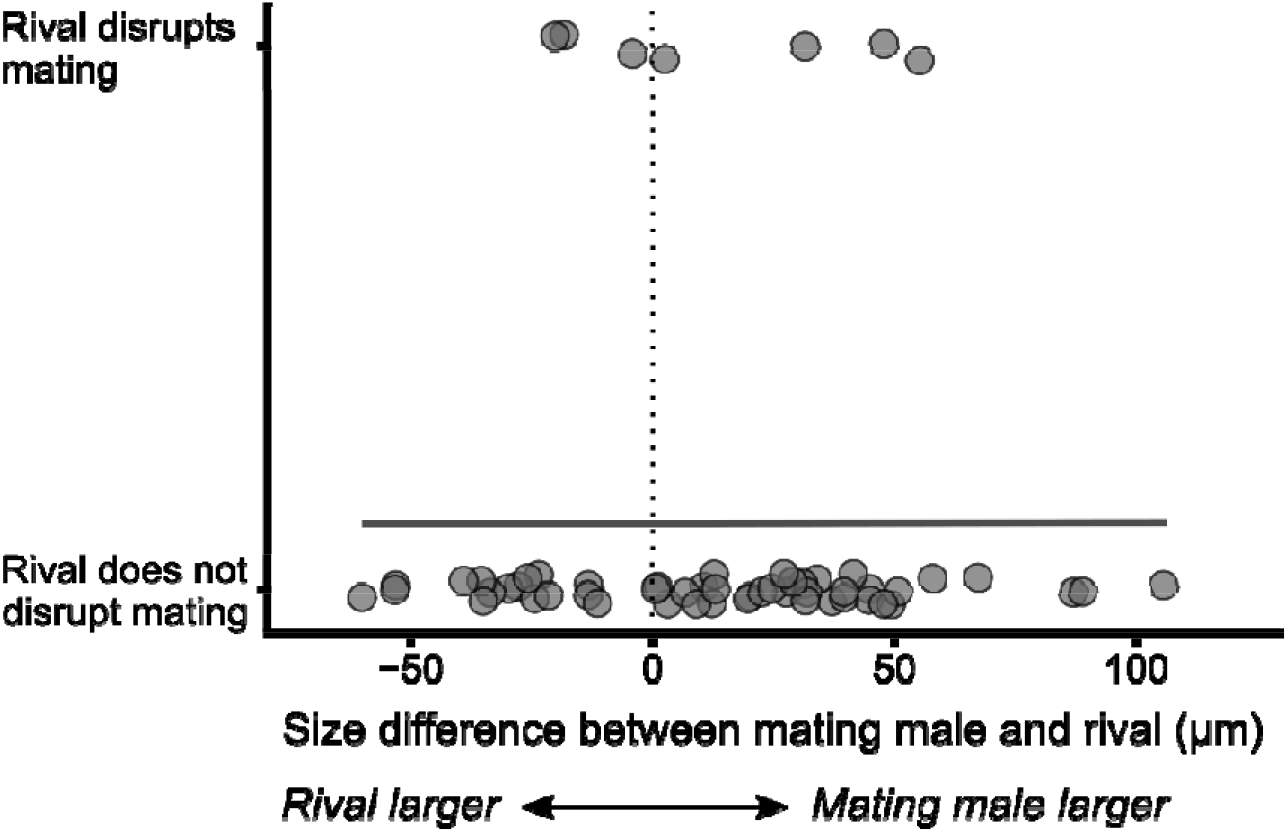
In fighter-fighter contests, the probability that an ongoing copulation is disrupted by the rival does not depend on relative body size. Copulation disruptions are rare. The black line represents the modelled probability of a disrupted copulation. The dotted vertical line denotes equal body size, at which the probability of disruption is estimated at 0.12 (95% CI: [0.06 – 0.25]).

Our fourth prediction was that copulations in the presence of a rival last longer if the rival is a scrambler rather than a fighter. Copulations by fighters lasted on average shorter when the rival was a fighter (median 13 minutes) than when the rival was a scrambler (median 18 minutes; Kruskal-Wallis χ^2^ = 6.07, df = 1, P = 0.014; Fig. 5). This was not due to interruptions of copulation in fighter-fighter contests, because also uninterrupted copulations by fighters lasted on average shorter when the rival male was a fighter, than when the rival male was a scrambler (Kruskal-Wallis χ^2^ = 4.97, df = 1, P = 0.026). Only three scramblers mated, so copulations by scramblers (with a rival fighter) were not considered in this analysis.

**Figure 5.**
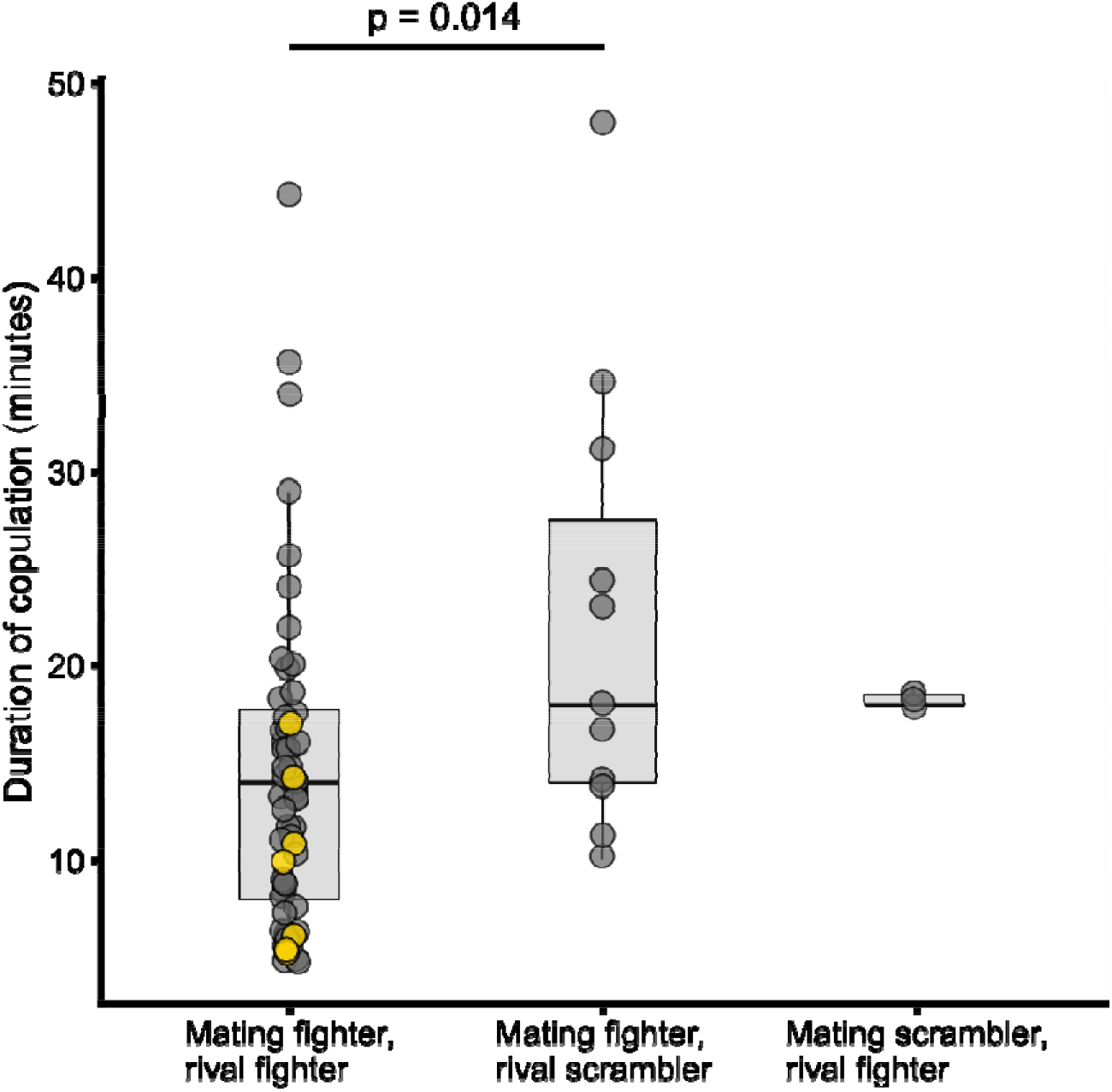
Copulations by fighter males last longer if the rival male is a scrambler, than if the rival is another fighter. This effect is not driven by disruption of copulations in fighter-fighter contests, as the effect persists when disrupted copulations (shown in yellow) are removed from the analysis.

## Discussion

We tested the hypothesis that condition-dependent male morph expression in *Rhizoglyphus robini* is associated with size-dependent alternative mating niches. Accordingly, only large males have access to the fighting niche, while smaller males resort to an alternative mating tactic. We found that males in fighter-fighter contests over a mating opportunity had a higher probability of mating if they were larger than their opponent, confirming that small fighters have disadvantage in the fighting niche (*prediction 1*). In fighter-scrambler contests, on the other hand, fighters had a higher probability of mating regardless of the body size difference, supporting that scramblers are not adapted to the fighting niche (*prediction 2*). We also found that copulations by fighters were occasionally disrupted by rival fighters, but only rarely, and apparently independent of the body size difference between the mating male and its rival (*prediction 3*). Copulations by fighters lasted shorter on average when the rival male was a fighter than when the rival was a scrambler (*prediction 4*), which could be due to the disturbing effect of the presence of another fighter. However, both uninterrupted and disrupted copulations lasted shorter in the presence of fighters, so shorter copulation durations were generally not attributable to aggressive physical disruptions of copulation by rival fighters. All in all, we conclude that the fighting niche in *R. robini* is particularly accessible for relatively large males who are able to win contests for mating opportunities. Small males, on the other hand, likely have to resort to alternative behaviours to mate. We found size-structured mating niche differentiation to be mainly due to size-dependent fighting success prior to copulation, and not due to size-dependent interruption of ongoing copulations. Size-dependent access to the fighting niche could play a role in explaining the evolution of condition-dependent male morph expression in *R. robini*.

Our results align well with evidence from horned beetles that larger males are generally more successful fighters than smaller males (Emlen 1997; Hongo 2007; McCullough and Simmons 2016; Mitchem et al. 2019), so that small males resort to sneaking behaviour to gain access to females, adaptively expressing a different, weaponless morphology (Moczek and Emlen 2000). We currently do not have evidence that *R. robini* scramblers are sneakers, but they have been shown to obtain higher fertilization success from a single copulation than fighters (Van den Beuken et al. 2019). Furthermore, scramblers move quicker than fighters and are better able to navigate spatially complex environments (Tomkins et al. 2011), which could benefit a hypothetical sneaking tactic. Hence, small males may be able to maximize fertilization success, given their poor fighting ability, by expressing the scrambler morph rather than the fighter morph, while large males benefit by expressing the fighter morph, given their high likelihood of winning fights. It should be noted that this scenario, in which fertilization success functions for scramblers and fighters cross each other over the body size gradient, has not been supported in a recent study. Specifically, Michalczyk et al. (2018) measured mass-dependent fertilization success of scramblers and fighters in the related mite *Sancassania berlesei*, which shows the same condition-dependent fighter-scrambler polyphenism as *R. robini*, but could not detect crossing fitness functions. On the one hand, this could indicate that reduced mating success in scramblers is not compensated by sneaking behaviour, and that condition-dependent scrambler expression might be adaptive for a different reason (e.g. to prevent immediate viability issues during metamorphosis (Smallegange et al. 2019)). On the other hand, Michalczyk et al (2018) may simply not have had enough statistical power to detect crossing fitness functions.

We found that fighter copulation time was shorter in the presence of a rival fighter than in the presence of a rival scrambler. This effect did not appear to be due to disruption of copulation by fighters. Social-context-dependent copulation duration in *R. robini* has been recorded previously (Skwierzyńska et al. 2018), where copulation duration increased slightly when males had previously experienced rivals from a different morph than themselves. This is consistent with our results, although we did not test whether scramblers increased copulation time in the presence of fighters rather than scramblers. Whether such behavioural plasticity is adaptive remains to be seen. It seems intuitive that males of both morphs would increase copulation duration in the absence of fighters, released from risk of attack. It is less intuitive that scramblers would increase copulation duration in the presence of fighters (Skwierzyńska et al. 2018).

In conclusion, we demonstrate that an aggressive male morph in *R. robini* increases mating success mainly for large males. Small males, on the other hand, are disadvantaged in aggressive contests, and benefit less by developing adaptations for fighting. The pattern that larger individuals are better able to secure mating opportunities than smaller individuals in intrasexual contests seems to be a general one (Parker 1974; Maynard Smith and Parker 1976; Lailvaux and Irschick 2006). In species with condition-dependent male polyphenism, this competitive asymmetry, and ensuing mating niche differentiation, likely contributes to maintenance of condition-dependent expression of alternative male morphs and mating tactics (Gross 1996; Oliveira et al. 2008). However, size-structured mating niches may not be the only factor involved in evolutionary maintenance of condition-dependent alternative morphs. Variation in growth (condition) impacts fitness in multiple ways beyond social status, and condition-dependent polyphenism may adaptively mitigate poor fitness prospects through e.g. reducing physiological constraints to rapid maturation, or maintaining viability by reducing allocation to expensive sexual traits (Smallegange et al. 2019). It remains to be seen whether small males actually maximize fertilization success with an alternative mating tactic such as sneaking behaviour, and are under selection for scrambler expression because of that particular effect. Nevertheless, size-structured mating niche differentiation, due to reduced fitness of a sexually selected phenotype in small males, undoubtedly facilitates the evolution of conditiondependent polyphenism.

